# MR relaxometry-based analysis of brain hemorrhages: an experimental study on a rabbit model

**DOI:** 10.1101/2021.01.12.426333

**Authors:** Francesca Del Signore, Massimo Vignoli, Leonardo Della Salda, Roberto Tamburro, Ilaria Cerasoli, Andrea Paolini, Mariarita Romanucci, Francesco de Pasquale

## Abstract

Magnetic Resonance Relaxometry is a quantitative MRI-based technique able to estimate tissue relaxation times T1 and T2. This approach allows increasing the MRI diagnostic accuracy mostly in case of brain neoplasia or neurodegenerative disorders in human medicine. However, few reports are available on the application of this technique in the clinical field of veterinary medicine. For this reason, in this work, we developed a relaxometry based approach on experimentally induced brain hemorrhages on rabbits. Specifically, the methodology is based on a hierarchical clustering procedure driven by the T1 relaxometry signals from a set of regions of interest selected on the T2 map. The approach is multivariate since it combines both T1 and T2 information and allows the diagnosis at the subject level by comparing “suspected” pathological regions with healthy homologous ones within the same brain.

To validate the proposed technique, the scanned brains underwent histopathological analyses to estimate the performance of the proposed classifier in terms of Receiver Operator Curve analyses. The results showed that, in terms of identification of the lesion and its contours, the proposed approach resulted accurate and outperformed the standard techniques based on T1w and T2w images. Finally, since the proposed protocol in terms of the adopted scanner, sequences, and analysis tools, is suitable for the clinical practice, it can be potentially validated through large-scale multi-center clinical studies.

## Introduction

Magnetic Resonance Imaging (MRI) is considered the gold standard imaging technique to investigate encephalic disorders in companion animals [1,2]. However, the images’ evaluation is typically qualitative since MRI provides signals only weighted on the actual tissue longitudinal (T1) and transverse (T2) relaxation times. To overcome this limitation and to obtain more objective information on the structure and function of the tissues under investigation, it has been developed a technique called MR-Relaxometry (MRR) able to provide maps of the actual T1 and T2 [3,4]. Although the intricate relationships linking the tissue microstructure to these relaxation times remain to be firmly established, their quantitative measurement can be informative of disease-related tissue change, developmental plasticity, and other biological processes. Therefore, MRR studies potentially offer a more detailed tissue characterization as compared to conventional ones [3,4].

In human medicine, the MRR role has been thoroughly investigated, for example in brain neoplasia, where this technique improved the lesion identification, especially when monitoring the chemotherapy response [5–9]. Many reports are also available on the application of this technique on epileptic patients, with the result of optimizing and enhancing lesion depiction [10,11,11–15]. In the case of neurodegenerative disorders, MRR resulted useful to improve the detection of early degeneration areas [16–18]. Additional examples and a general review of this technique can be found in [3,4].

As far as it concerns Relaxometry applications on animal models, most studies have been performed on rats, mainly on neoplasia [6,19,20]. Further, MRR has been useful in characterizing the hippocampal involvement in epileptic patients [21] and to detect spontaneous brain hemorrhages [22]. However, most of the animal studies lack an accurate histopathological validation and are typically performed with high field scanners characterized by a high spatial and signal-to-noise-ratio (SNR) [21]. On the one hand, this allows us to achieve a higher spatial resolution and signal-to-noise ratio (SNR). On the other hand, translating such protocols in the clinical field might not be straightforward since nowadays, mainly low field scanners, characterized by a low spatial resolution and SNR, are available in veterinary facilities [23]. Typically, the diagnosis is based on standard T1w/T2w images and small lesions can be missed. Further, it often strongly relies on the radiologist’s experience, e.g. in case of subtle lesions, hepatic encephalopathy, distemper encephalitis, or small hemorrhages [24,25]. To overcome these limitations, in the clinical field, an advanced MRR based tool, as long as compatible with a low field scanner equipment, could be helpful, as shown in [22]. In this preliminary study, we showed the potentiality of a relaxometry protocol implemented on a 0.25 T scanner and tested on phantom and real data from a healthy dog [22]. Here, we built upon those results to focus on experimentally induced hemorrhages on rabbit brains. This model allows to work with brains whose size is similar to canine and feline species so that the developed protocol might be translated to these species. Further, smaller brain animals were excluded due to the resulting prohibitive SNR of the adopted MR scanner.

The developed approach can be summarized as follows. First, a set of regions of interest were selected based on a T2 map. Second, a hierarchical clustering, driven by the T1 relaxation patterns of the selected regions, compared the selected regions with the healthy homologous ones and classified the tissues under investigation. Finally, the performance of the proposed classifier was assessed through the comparison with histopathological examinations of the brains under investigation.

## Materials and methods

### Cerebral hemorrhage model in rabbits

Sixteen New Zealand White male rabbits (*O. Cuniculus*) (3.2 ± 0.5 kg) were included in this study, that has been carried out in strict accordance with the recommendation from national committee for animal welfare, with the procol N° 726/2019-PR; all the subjects received continuous proper care according to national guidelines under the supervision of trained personal.

Eight hours before the procedure, each rabbit received meloxicam (1,5 mg kg −1) (Metacam, 1,5 mg/ml, Boehringer Ingelheim Div. Veter) intramuscular (IM) to obtain multimodal analgesia integrated with morphine (1 mg kg −1) (morphine chloride, 10 mg/mL, S.A.L.F. S.p.a) and midazolam (0,5 mg kg −1) (Dormicum^®^, 5 mg/ml, Roche Pharma, Switzerland) for IM premedication before the start of the procedure. Sixty minutes before applying auricular venous and arterial catheter (22-24 G 2,5 cm) to properly administer fluids and drugs as needed, and measure blood pressure respectively, each rabbit received topic administration of EMLA cream 5% ^®^ (Aspen Pharma Trading Limited). Induction has been performed with alfaxalone (1-4 mg kg −1) (Alfaxan^®^, 10 mg/ml, Jurox, UK) intravenous (IV) and general anesthesia was maintained with isoflurane in oxygen 100% after applying a cuffed orotracheal tube (2-3-5 mm ID) under endoscopic guidance.

The rabbit cerebral hemorrhaged model was established by autologous blood injection method, with 0.15 ml of blood extracted from the auricular vein. The rabbit was placed in sternal recumbency; a 1.5 cm midline sagittal skin incision was performed with a #10 scalpel blade and the subcutaneous tissue was dissected to visualize the brain “cross stitch” intersection that was set as the starting point. A 2.0 mm bone defect was manually performed on the left calvarium with a Synthes-Colibri 2 surgical drill using a Synthes drill bit diameter 2.0 mm. The drilling starting point was 3-5 mm to the left of the coronal suture and caudal to the sagittal suture [26] (Fig 1). The drilling was perpendicularly oriented to the parietal bone surface. A stopper was applied 4 mm below the tip of the drill bit to avoid any potential brain damage. Then the blood was manually injected into the left hemisphere of each subject with a 1 ml syringe and a 25 G-1.6 cm needle perpendicularly oriented to the parietal bone surface to compare the two hemispheres in the same acquisition.

**Fig 1.**
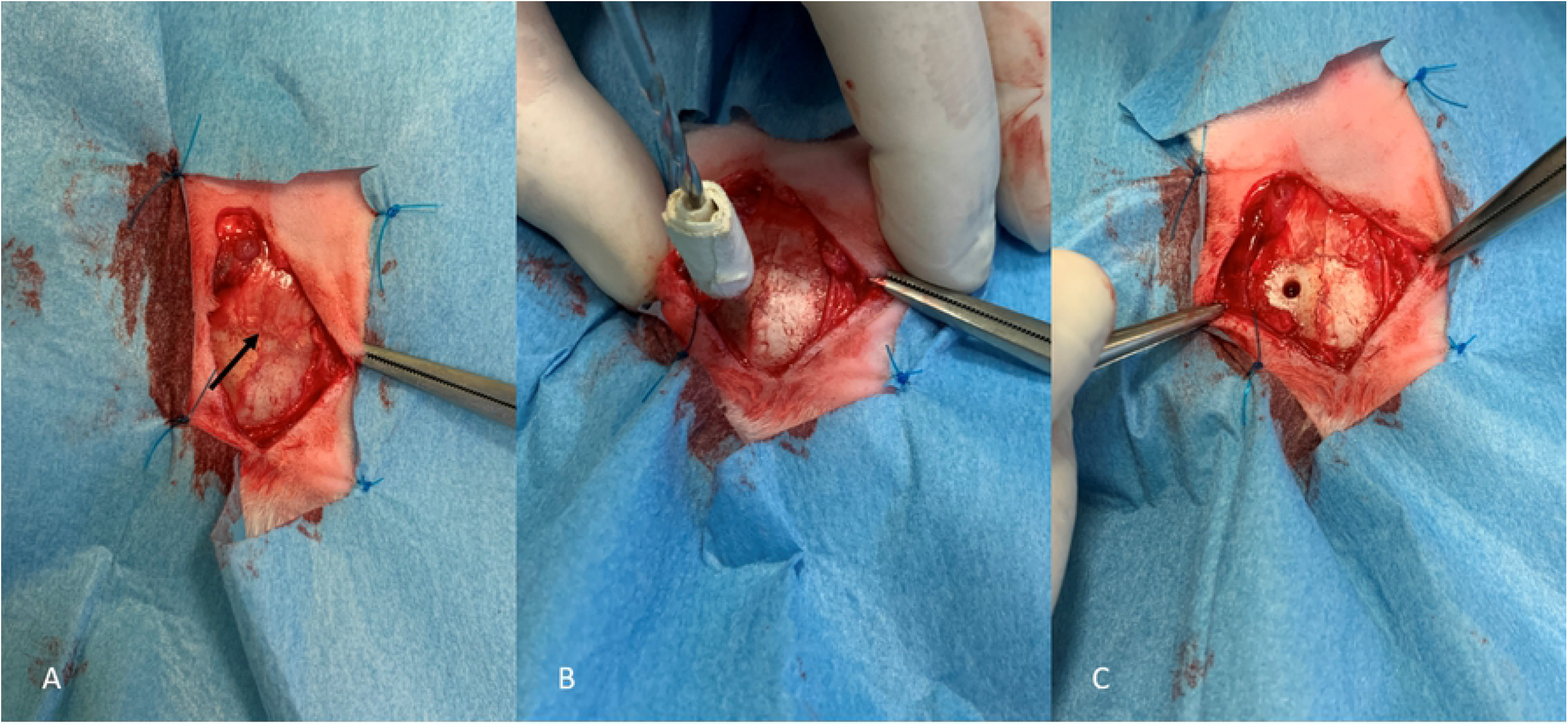
Technical procedure to induce hemorrhages. A) The brain “cross stitch” used as an anatomic landmark (black arrow). B) The 2-mm diameter hole performed with a surgical drill 3-5 mm caudally and on the left of the coronal suture. C) The final location to inject the autologous blood and induce the cerebral hemorrhage.

Then the rabbit was carefully moved to the MR scanner. After the MR acquisitions, each subject has been euthanized according to national guidelines, and brains were promptly fixed in 10% neutral buffered formalin for histopathological investigations (see below).

### MRI acquisition and relaxometry protocol

Thirteen out of sixteen subjects were included in the study since three rabbits died. MR data were acquired using an Esaote Vetscan Grande scanner operating at 0.25 T equipped with a Coil 4 (Esaote S.PA, Genova, Italy). The conventional data consisted of Spin Echo (SE) T1w and Fast SE T2w sequences on transverse and sagittal planes, see Table 1 for the adopted parameters.

**Table 1.** MRI acquisition parameters

|  | **T1w** | **T2w** | **T2w** |
| --- | --- | --- | --- |
|  | **Transverse** | **Transverse** | **Sagittal** |
| **TR** | 400 ms | 2660 ms | 2170 ms |
| **TE** | 18 ms | 100 ms | 100 ms |
| **Slice thickness** | 3 mm | 3 mm | 3 mm |
| **Gap** | 0 mm | 0 mm | 0 mm |

Settings for T1w -T2 w transverse and T2 w sagittal sequences.

Based on these acquired images, a single transverse brain slice was selected to perform both T1 and T2 relaxometry. This slice was selected to contain the suspected localization of the lesion in the transverse plane. When the lesion was not clearly identifiable in this plane, sagittal sequences were used to localize the lesion.

T1 Relaxometry data were acquired through an ad-hoc protocol based on repeated acquisitions of SE T1w images corresponding to a variable TR, namely TR = [50 120 200 300 400 500 600 650 750 850 900 950 1000] ms. This set of values has been optimized in a previous study [22]. As far as it regards T2 relaxometry data these were obtained by acquiring FSE T2w images repeatedly acquired with a variable TE, namely TE = [28 75 136] ms.

To account for small head movements during the acquisition, all the acquired images were coregistered to the T1w ones, through an affine transformation using a standard approach, see for example [27]. For this reason, in what follows all the acquired images and related results, such as T1/T2 maps and clustering results, will result aligned in the same subject space. The complete acquisition procedure lasted about 45 minutes on every subject.

### Analysis pipeline

The proposed approach consists of a multivariate classification procedure that combines both the information from T1 and T2 relaxation times. The analysis pipeline is schematized in S1 Fig.

#### Estimation of T1 and T2 maps

To estimate T1/T2 maps the acquired MR signals were fitted through a two-parameter model using an unconstrained minimization algorithm based on a derivative-free method [28,29]. Then, for each patient, from the obtained T2 maps a set of *n* voxels, representing ROIs of suspected lesions, were manually selected by an expert radiologist (S1A Fig). The value of *n* was variable depending on the extension of the suspected lesion. Then, based on the location of the brain midline, the algorithm, in the contralateral hemisphere, automatically extracted additional *n* voxels corresponding regions homologous to the ones selected (S1A Fig). The set of 2*n* voxels will represent the input of the classification algorithm. This allowed us to compare the selected ROIs with their homolog ones to obtain subject-specific results avoiding the normalization into an average-based brain template.

#### Hierarchical clustering

To detect the hemorrhagic vs healthy tissues a hierarchical clustering, driven by the T1 relaxometry patterns (S1B Fig), was employed (S1C Fig). Other possible clustering techniques could have been adopted, but the hierarchical clustering has the advantage, e.g. as compared to K-means clustering, that the number of classes does not need to be specified in advance and the progression of the clustering can be evaluated at different levels.

The basic idea is that hemorrhagic tissues will exhibit a different pattern of relaxometry as compared to healthy ones. At this stage, we decided to use the original relaxometry pattern for every voxel and not the relaxation times at the previous steps. This makes the classification more robust since it depends on 13 points (the number of TR adopted), and not just 1 as in the case of relaxation time, thus describing more accurately the return of the MR signal to the baseline.

Specifically, the hierarchical clustering was run with the Euclidean distance and an unweighted average as linkage criterion. The final output of the hierarchical clustering is a dendrogram showing the hierarchical evolution from a set of completely separated observations to the final single cluster, (S1C Fig). At this stage, to choose the optimal number of clusters, for every output, we computed the cluster silhouette [30] Of note, for all the considered rabbits in this study, we obtained the highest silhouette corresponding to two classes, i.e. a binary classification. Thus, in what follows, the cluster of voxels whose majority was in the healthy hemisphere was labeled as “healthy” (H) while the other cluster as “pathological” (P). Of note, the healthy hemisphere will always be the right one since we induced hemorrhages only in the left hemisphere.

All the reported analyses were performed through in-house developed codes in MATLAB (MATLAB and Statistics Toolbox Release 2015b, The MathWorks, Inc., Natick, Massachusetts, USA).

#### Histopathological examination

Brains were cut with a small guillotine specifically prepared for the experiment (see Fig 2). Each brain has been positioned on a home-made wooden wedge to respect the same brain angle inclination measured from MRI images during the scan protocol; this aspect has been carefully considered to orient the cut parallel to the induced lesion (Fig 2).

**Fig 2.**
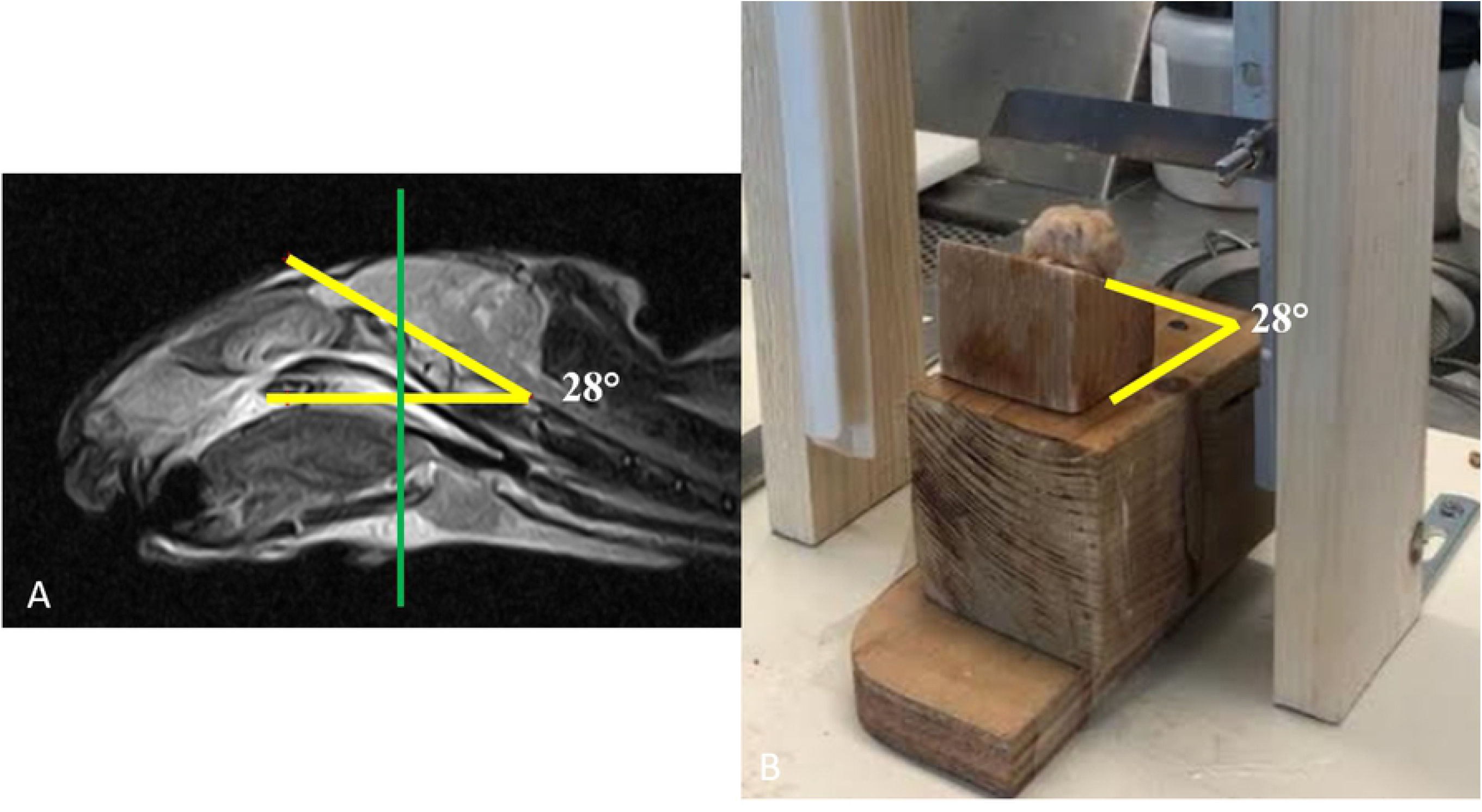
Operative procedure for brain cuts. A) The inclination of the brain obtained from the T2w sagittal images. B) The same inclination as in A) has been reproduced with wooden wedges to be used with a small guillotine specifically prepared for the experiment.

Transverse serial sections of all fixed brains were also obtained and photographed. Then, formalin-fixed paraffin-embedded brain sections (i.e, tissue blocks from all induced hematoma areas, as well as from ipsilateral and/or contralateral nervous tissues, particularly frontal Cerebral Cortex, Corpus Callosum, and Caudate Nucleus) were routinely processed for histology and stained with hematoxylin-eosin (H&E). The acquisition of the histological images and the evaluation of the margins of the lesions were carried out using the image analysis software LAS X Measurements (Leica microsystem).

### Validation of the proposed approach with histopathological analyses

To validate the proposed approach, the classification results were compared to the histopathological findings as follows.

The classifier performance was assessed through a Receiver Operator Curve analyses where the ‘ground truth’ was represented by the histopathological results (S1D Fig) First, based on the histological analyses, the contours of the tissues involved with the hemorrhages were extracted. In what follows, we will denote this contour as the “true lesion contour”. Then, these images were manually co-registered to the anatomical MRI images by an expert radiologist. In this way, for every rabbit, we could directly compare the classification with the ground truth. This allowed to compute, for every lesion, the true positives (TP), as the number of voxels correctly classified as pathological (red voxels in the figures), i.e. voxels falling within the “true lesion contour”; the false negatives (FN), as the number of voxels classified as healthy falling inside the “true lesion contour”; the false positives (FP), as the number of voxels classified as pathological that fell outside the “true lesion contour” and the true negatives (TN) as the number of voxels classified as healthy (yellow in the figures) falling outside the “true lesion contour”. This allows computing the accuracy, true positive rate (TPR), and false-positive rate (FPR) according to the available literature [31]. Based on these parameters, we computed the ROC and the corresponding area under the curve (AUC) that can be interpreted as follows:

AUC = 0.5 the test is not informative;
0.5 <AUC ≤0.7 the test is not accurate;
0.7 <AUC ≤0.9 the test is moderately accurate;
0.9 <AUC <1.0 the test is highly accurate;
AUC = 1 perfect test [32].

ROC analyses were performed using an in-house developed code in MATLAB (MATLAB and Statistics Toolbox Release 2015b, The MathWorks, Inc., Natick, Massachusetts, USA).

## Results

### Conventional MRI vs MR Relaxometry

First, the conventional MR protocol was tested to identify the induced lesions. Based on T1w images, a board-certified radiologist judged the lesion as ‘not visible’ in 46% of the subjects and ‘barely visible’ in the remaining 54%. As an example, in Figs 3–5A the T1w images of three representative patients are shown. It can be noted that with that the typical settings of T1w sequences, the induced hemorrhages were not clearly identifiable in two of the reported cases (Figs 3 and 4A), while it was barely visible in the last one (Fig 5A). The visualization of the lesions improved when T2w images were considered (Fig 3–5B): the rate of lesions classified as ‘not visible’ dropped to 23% while the proportion of ‘clearly visible’ and ‘barely visible’ raised to 39% and 38%, respectively (see Figs 3–5B-white arrows). As it can be seen in the reported examples, lesions were characterized by ill-defined patterns whose extension varied throughout the sample. In 22% of the sample, the lesions were localized in the left frontal lobe (Fig 4), and in the remaining 78%, the lesions were localized in the left thalamus (Figs 3 and 5).

**Fig 3.**
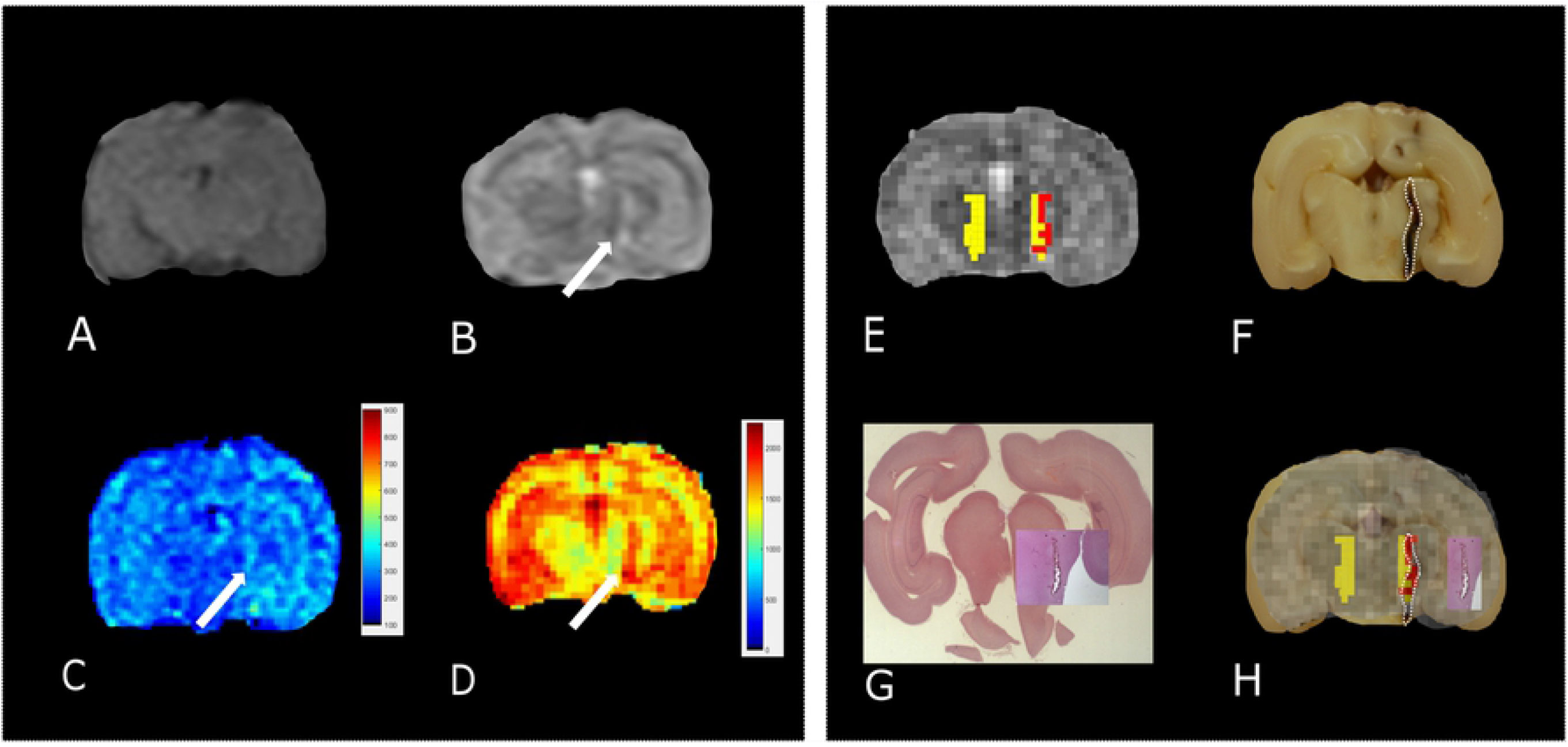
Standard vs MRR analyses and histopathological validation. Example 1. A) T1w transverse image of the brain; the lesion has been classified as ‘not visible’. B) T2w transverse image of the brain; the lesion appears ill-defined, hyperintense, and linearly shaped in the left thalamus (white arrow). C-D) T1 and T2 maps, respectively. A hyperintense linearly shaped area is visible in both maps (white arrow). E) The output of the hierarchical cluster overlaid on the T2 map. Voxels classified as “pathological” and “healthy” parenchyma are reported in red and yellow, respectively. F) The results of gross anatomy. From the photograph of the brain section, the contour of the hemorrhagic lesion characterized by a linear shape that mainly involves the left thalamus (white arrow) has been delineated (white dotted line). G) Histopathology results. The overall brain section obtained from the microscope with the hemorrhagic lesion shown in greater detail superimposed on the overall section. H) The MRR classification compared to both gross anatomy and histopathology: voxels classified as pathological (red) fall inside the estimated true lesion contour.

**Fig 4.**
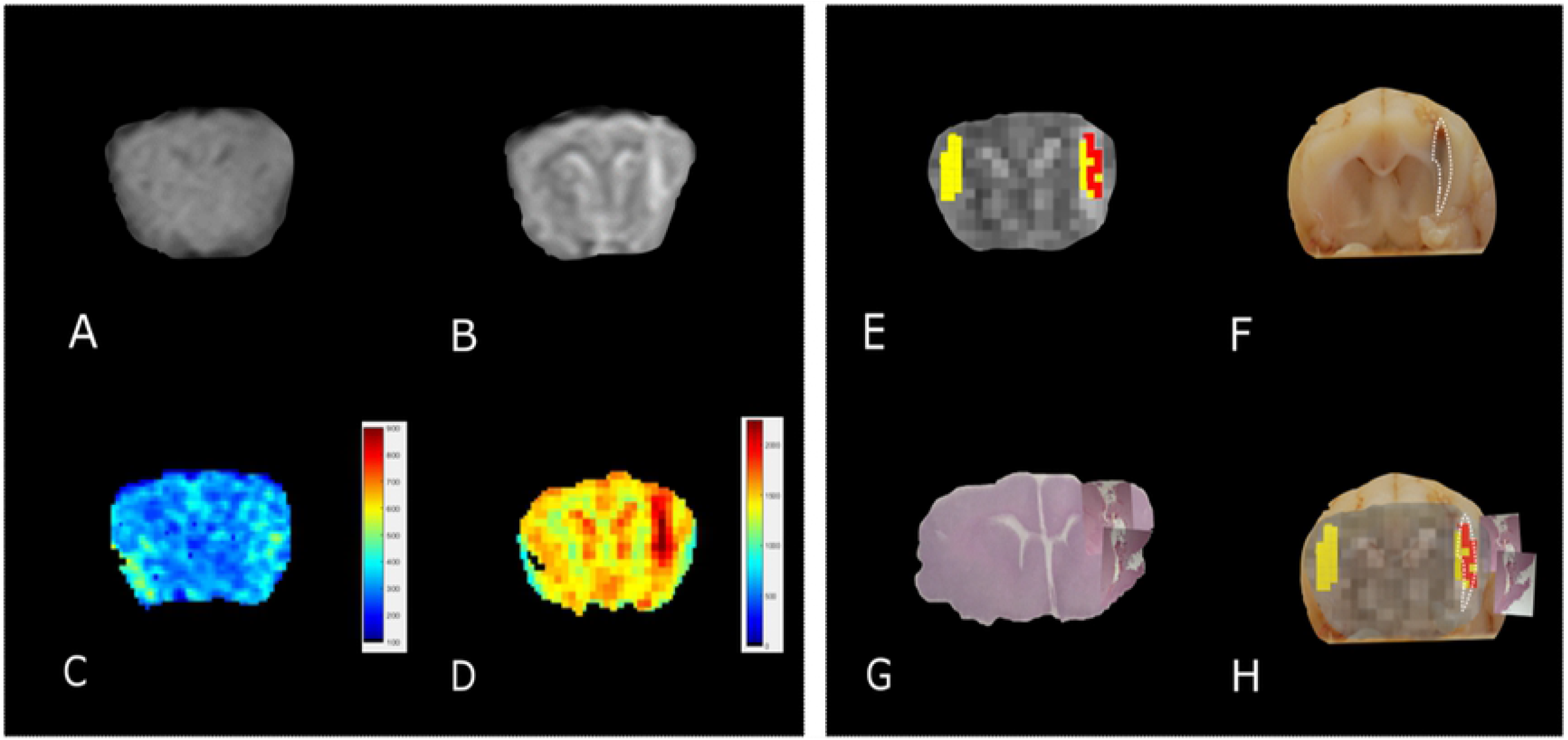
Standard vs MRR analyses and histopathological validation. Example 2. A) T1w transverse image of the brain; the lesion has been judged ‘not visible’ by a board certified. B) T2w transverse image of the brain. Also in this case the lesion appears ill-defined, hyperintense and linearly shaped in the left thalamus (white arrow) C-D) T1 and T2 maps, respectively. A hyperintense linear-shaped area is visible in both maps (white arrow). E) The output of the hierarchical cluster overlaid on the T2 map. Voxels classified as “pathological” and “healthy” parenchyma are reported in red and yellow, respectively. F) The results of gross anatomy. From the photograph of the brain section, the contour of the hemorrhagic lesion characterized by a linear shape that involves the left frontal lobe (white arrow) has been delineated (white dotted line). G) Histopathology results. The overall brain section obtained from the microscope with the hemorrhagic lesion shown in greater detail superimposed on the overall section is reported. H) A good agreement can be noted between the MRR classification and the lesion contour estimated from both gross anatomy and histopathology.

**Fig 5.**
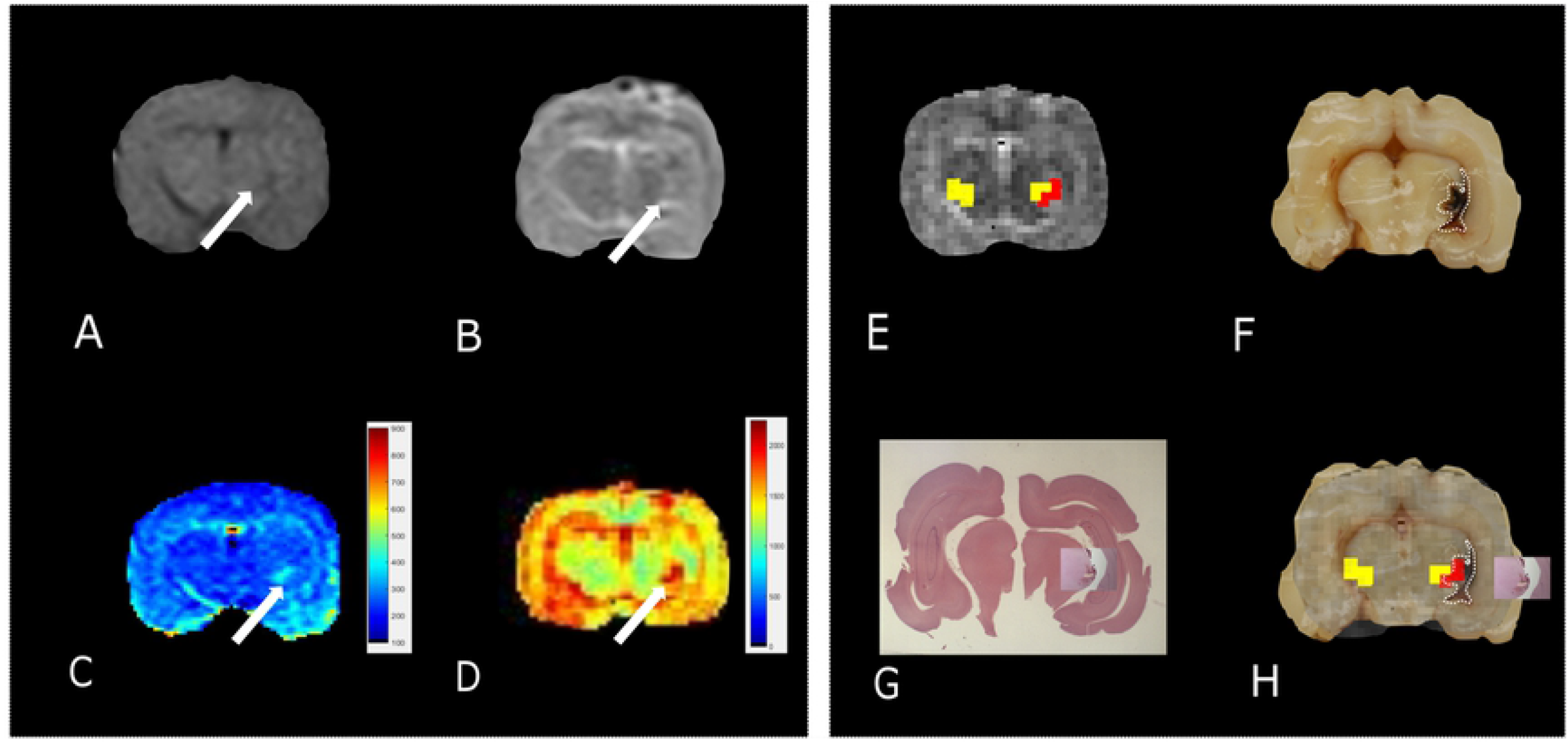
Standard vs MRR analyses and histopathological validation. Example 3. A) T1w transverse image of the brain; the lesion appears ill defined, hypointense, and irregular in the left thalamus (white arrow). B) T2w transverse image of the brain. As in T1w, the lesion in the left thalamus appears ill-defined, irregular but hyperintense (white arrow). C-D) T1 and T2 maps, respectively. A hyperintense and irregularly shaped area is visible in both maps (white arrow). E) The output of the hierarchical cluster overlaid on the T2 map. Voxels classified as “pathological” and “healthy” parenchyma are reported in red and yellow, respectively. F) The results of gross anatomy. From the photograph of the brain section, the contour of the hemorrhagic lesion that appears irregular in shape and involves the left thalamus (white arrow) has been delineated (white dotted line). G) Histopathology results. The overall brain section obtained from the microscope with the hemorrhagic lesion shown in greater detail superimposed on the overall section. H) The hemorrhagic lesion found by the MRR approach (red voxels) is in agreement with the gross anatomy: voxels classified as pathological fall inside the true lesion contour. It can be noted that in panel F) the hemorrhagic area appears more extended than the lesion histologically evident (G). However, this can be due to postmortem blood spread in the space between the left hemisphere and the ipsilateral thalamus. The contour of the hyperintense area in both T1 and T2 maps is in very good agreement with the hemorrhagic area evident from the histopathological results where the blood spread disappeared after the formalin fixation.

Then, the T1 and T2 maps were estimated. In this case, a further improvement was obtained in terms of lesion identification: lesions were judged as ‘clearly visible’ in most of the sample, namely 85% for T1 maps (see Figs 3–5C) and 93% for T2 maps (see Figs 3–5D). In the remaining 5% and 7% for T1 and T2 maps respectively the lesion was not clearly detectable. In general, as it can be seen in Figs 3–5CD (white arrows), hyperintense lesions were observed with punctiform, linear, or irregular shape whose size varied across subjects.

Eventually, hierarchical clustering was performed to classify brain tissues. As described in Section 2, a set of candidate voxels were selected based on T2 maps. As an example, we report in S1 Fig (see supplementary material) voxels selected (black dots) and their homologous ones (red dots) for the data of Fig 3. Based on their T1-relaxation signals (see S1B Fig), the hierarchical clustering was run. The final classification of healthy (H-yellow) and pathological (P-red) voxels is reported in Figs 3–5E overlaid on the T2 map (grayscale). Interestingly, in all reported cases, within the initially selected area, the hierarchical clustering identified a subset of ‘pathological’ voxels. Theoretically, since lesions were induced only in the left hemisphere, members of the P class were not expected in the right hemisphere. This applied to the reported cases of Figs 3–5.

However, there have been cases where classification partially failed and labeled as pathological also voxels in the healthy hemisphere, see for example Fig 6.

**Fig 6.**
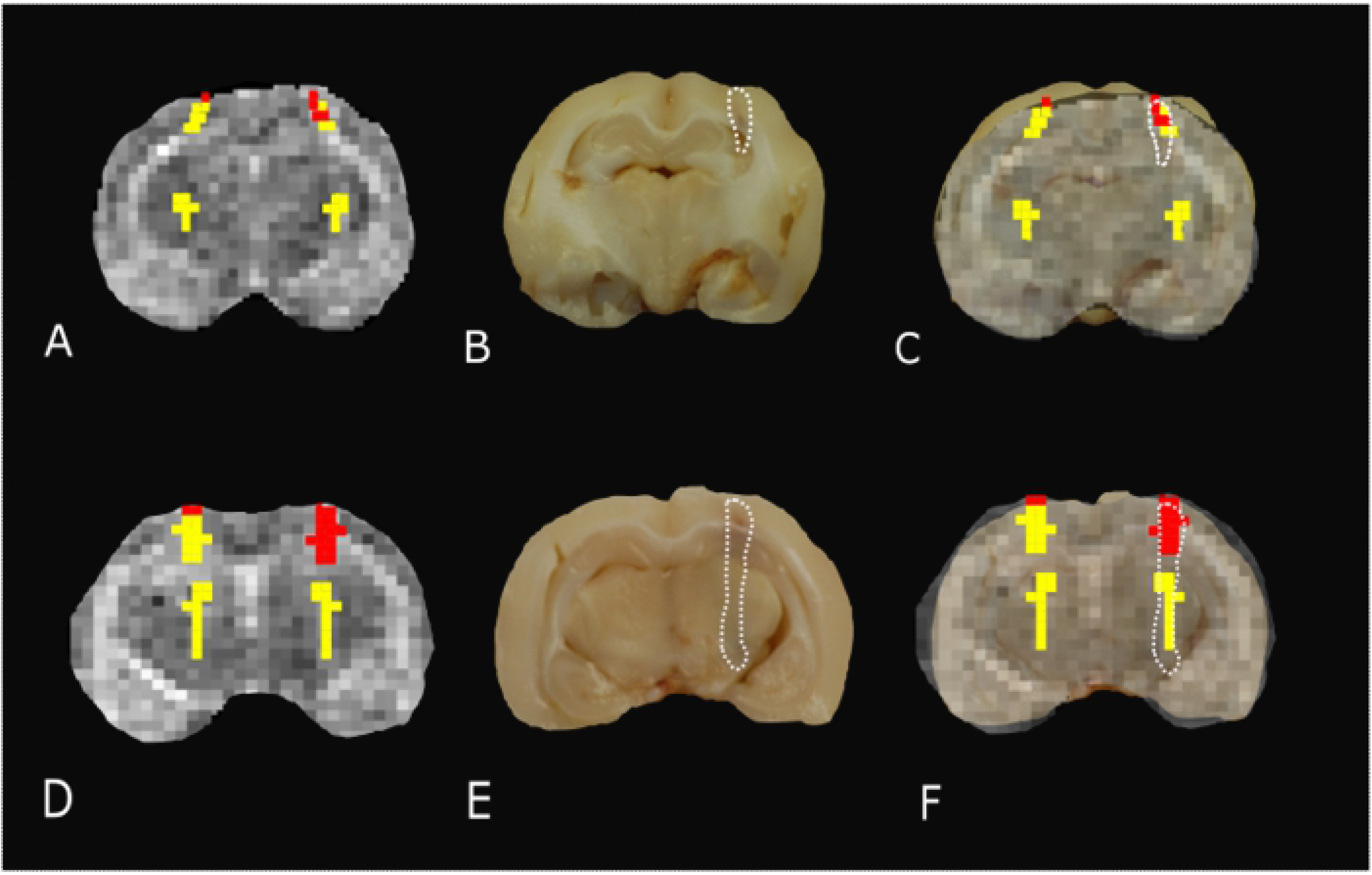
Misclassification results. A, D) The results of MRR classification do not show a clear distinction between the two groups of pathological vs healthy parenchyma, since pathological tissue was found in the healthy hemisphere. B, E) The hemorrhagic lesion contour (white dashed line) estimated from the gross anatomy. C-F) The comparison between the MRR classification and the true lesion contour shows that in these two cases the performance of the classifier is poor since both healthy (yellow) and pathological (red) voxels fall inside the lesion contour.

This aspect may be justified due to blood heparinization that may have led to a lack of a clear division between the hemorrhage and the brain parenchyma, see Discussion.

These findings show that it is fundamental to validate accurately the proposed approach. To this aim, we assessed the classification performance by comparing these results with the histopathological assessment.

### Histopathology assessment

The induced hemorrhages varied in size and shape, and a superficial, subdural, or subarachnoid collection of blood was also usually observed. Most frequently affected areas were represented by the frontal cerebral cortex (gray and white matter), cingulate gyrus, and caudate nucleus.

Histologic evaluation was performed to identify the pattern of hemorrhage and perihaemorrhage tissue changes. Serial sections allowed to observe the extension of the hemorrhagic area caused by the needle-induced direct trauma and the aspect of perilesional tissues, not affected by the blood accumulation. In all the brains, a hemorrhagic lesion was detected with a pattern characterized by a moderate variation of intensity from case to case (Fig 7). The injected blood caused disruption of the surrounding neural tissue, termed mass effect (Fig 7A, B) Fragmented nuclear debris was observed in damaged perihematomal tissue, in association with the presence of perihematomal edema, frequently characterized by gray matter vacuolization, neuronal perinuclear halos (Fig. 7C black arrow; 7D black arrow), and multifocal, mild dilation of perivascular (Virchow-Robin) spaces. Red blood cells within the wall of small vessels (intramural erythrocytes) and scattered swollen endothelial cells were also seen (Fig 7D white arrow). Also, multifocal cell shrinkage, with the presence of scattered hypoxic-ischemic neurons was detected in almost all cases (Fig 7D arrowhead; 7E arrow). For comparison, neural tissues distant from the hematoma in the same section, as well as the corresponding contralateral areas in the unaffected cerebral hemisphere were examined, and no significant alterations were recorded, thus allowing to exclude the occurrence of histologic artifacts due to brain fixation or manipulation (Fig 7F).

**Fig 7.**
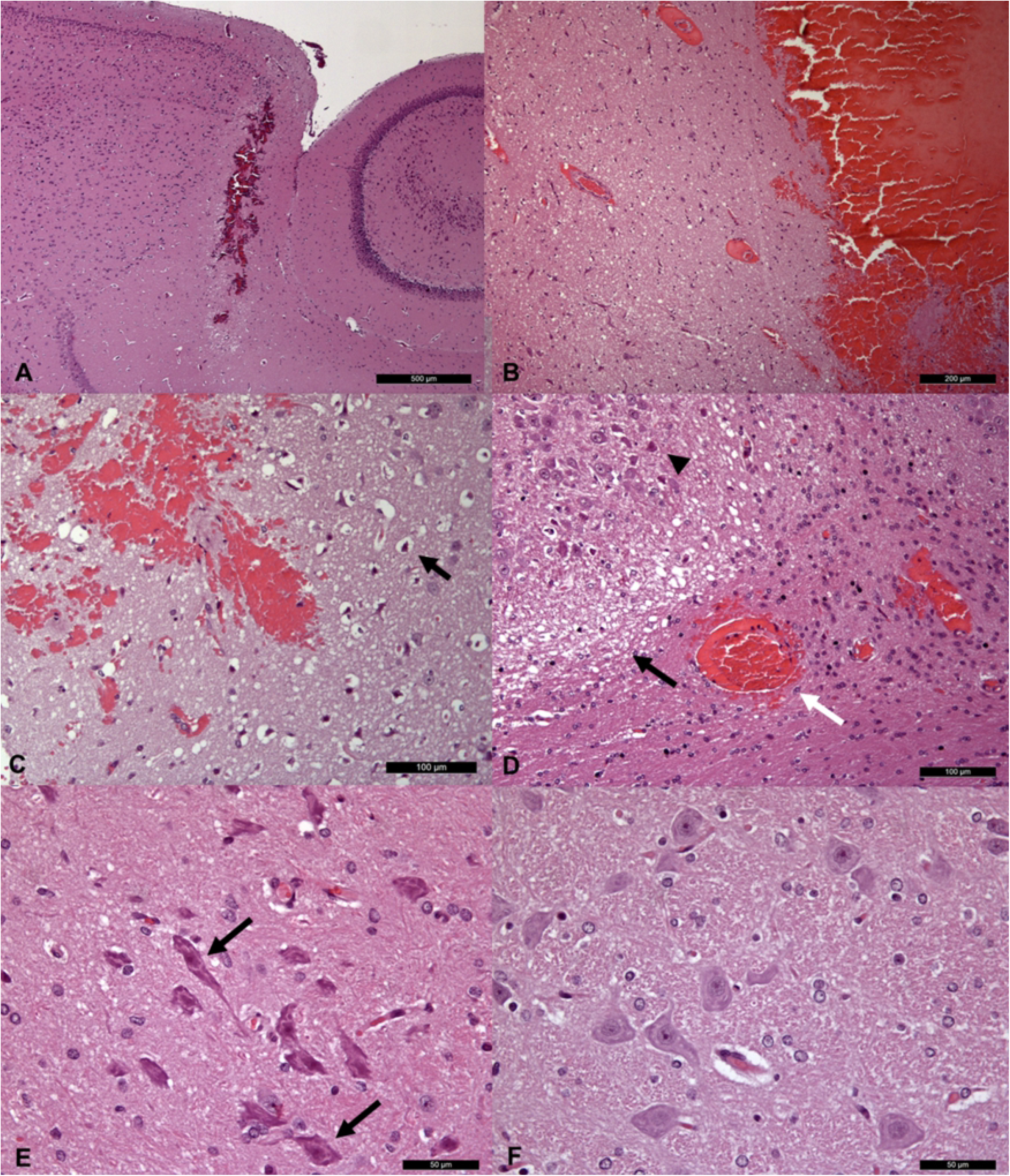
Histopathological sections. A-B) Low (A) and high (B) magnification of brain sections passing through the needle path, showing a linear hemorrhagic area. C-D) Histological images of peri-hemorrhage neural tissue, showing variably intense, (C) gray (perinuclear halos; arrow) and (D) white matter edema (black arrow) and multifocal neuronal shrinkage (arrowhead). Red blood cells within the wall of a small vessel are also visible (white arrow). E) Higher magnification of multifocal neuronal shrinkage (arrows) in peri-hemorrhagic tissue. F) Histologically normal neurons in the contralateral area of the unaffected cerebral hemisphere.

The above aspects evidence the structural difference between the brain hemisphere with induced hemorrhage and the healthy one, thus confirming that the hyperintense areas on T1 and T2 maps were not artifactual in nature and correspond to real lesions.

### Validation of the proposed approach

To validate the proposed approach, we compared, through a ROC analysis, the MRR results with the histopathological ones described above. For every patient, the lesion contours were manually delineated from both gross anatomy and histopathological images. In Figs 3–5F, we show the lesion contour (dashed line) extracted from a photograph of the section of the brain immediately after the extraction, typically within 10-15 minutes after the end of the MRI session. This section was selected as explained in Section 2, to match the MRI transverse plane where the relaxometry study was performed. Furthermore, the lesion contour was also obtained from the gross anatomy sections and microscopic analyses performed at 48 hours after the brain was kept in formalin. The macroscopic image of the brain section with an inset from the microscopic analyses of the hemorrhages is reported in Figs 3–5G. As it can be noted in Figs 3–5H, where the classification and anatomy results are superimposed, a good agreement between the gross anatomy (dashed) and MRR classified voxels (red) was obtained. However, as far as it regards the cases shown in Fig 6, the classification seems to match only the initial part of the lesion (see Fig 6C) or half of it (Fig 6F). Now, to quantitatively validate the MRR approach, (see Section 2) the TPR and FPR for each lesion were computed. The results reported in Table 2 seem promising since the mean TPR was 0.76, the mean FPR was 0.13 leading to an overall accuracy of 0.83.

**Table 2.** ROC analyses.

| **RABBIT** | **TRUE**<br>**POSITIVE**<br>**RATE (TPR)** | **FALSE**<br>**POSITIVE**<br>**RATE (FPR)** | **ACCURACY** |
| --- | --- | --- | --- |
| 1 | 0.87 | 0.34 | 0.77 |
| 2 | 0.62 | 0.18 | 0.74 |
| 3 | 1 | 0 | 1 |
| 4 | 0.7 | 0.3 | 0.7 |
| 5 | 0.67 | 0.05 | 0.89 |
| 6 | 0.67 | 0.2 | 0.74 |
| 7 | 1 | 0.15 | 0.92 |
| 8 | 0.34 | 0 | 0.67 |
| 9 | 1 | 0 | 1 |
| 10 | 0.95 | 0.17 | 0.88 |
| 11 | 1 | 0 | 1 |
| 12 | 0.3 | 0 | 0.65 |
| 13 | 0.91 | 0 | 0.96 |
| **MEAN** | **0.76** | **0.12** | **0.83** |

TPR, FPR, and accuracy are reported for each rabbit.

In Fig 8 the ROC curve is reported (the point 1 and 0 are reported only for clarity purpose to show to reference bisecting line). As it can be noted, the classification performance is well beyond the chance (0.5, dashed line) level, being the area under the curve AUC = 0.87. This value, compared to the thresholds reported in Section 2, places the performance of the classification close to the edge of “highly accurate” level (AUC = 0.9).

**Fig 8.**
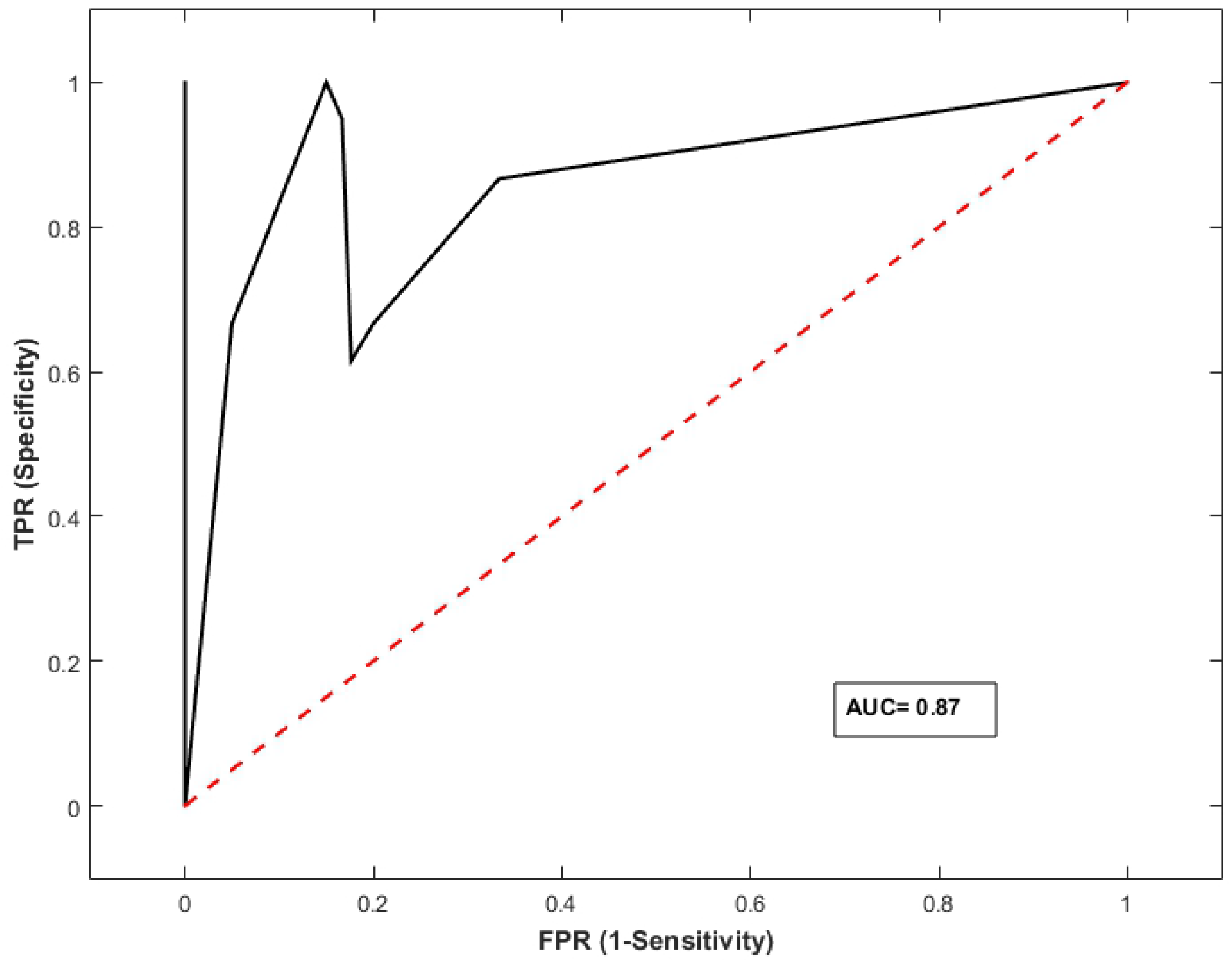
ROC analysis. The ROC curve shows that the classifier performance (black line) is beyond the chance level (red line). The area under the curve (AUC=0.87) compared to the thresholds reported in Section 2, places the performance of the classification close to the “highly accurate” level (AUC = 0.9).

## Discussion

In this study, we proposed a multivariate MRR approach to identify subtle brain hemorrhages on a rabbit model. The developed technique, which supports a diagnosis at the subject level, has been compared to conventional MRI and validated through histopathological examinations. The proposed approach, being optimized for a low field scanner, routinely available in the veterinary facilities, has the advantage of a potential direct translation into the clinical practice.

In general, the MRI sensitivity for detecting subtle lesions such as edema or hemorrhages increases with the magnetic field. Thus, with low field scanners typically used in veterinary practice, these hemorrhagic diseases are likely to be underdiagnosed [25]. For this reason, we tested our approach on induced hemorrhagic lesions obtained by injecting a minimal amount of blood to create a reproducible and lateralized lesion. This allowed to compare the injured and healthy parenchyma within a single MR scan. This approach is in line with previous works in the literature, for example in [33] the authors injected autologous blood in the right hemisphere of rat brains to evaluate the effects of experimental intracerebral hemorrhage on brain tissue injury and recovery. They reported that MR relaxometry can be promising to observe the lesion progression. Our choice to focus on T1 and T2 Relaxometry, instead of T2*, shown also reliable in visualizing hemorrhage [34], is also in line with previous MRR studies on this kind of lesions [22,26,33]. In fact, to setup T2* relaxometry protocols can be very challenging with a low field clinical scanner, and thus T1/T2 MRR has the advantage of being more easily implemented in the future by veterinary practitioners on a large scale. The proposed approach is multivariate, i.e. it combined both T1 and T2 information: the highest contrast in the T2 map was exploited to select the initial ROIs, while the hierarchical clustering was driven by the T1-based signal recovery. In fact, in terms of qualitative evaluation, in our data, the lesions resulted always more evident in T2 maps than in T1 ones. Moreover, the adoption of the T1 signal to drive the classification is in line with the literature where T1 relaxometry has been proved to be useful to assess brain parenchyma even with low filed scanners [22,35]. Nevertheless, this choice leads to a higher acquisition time, which is one of the main drawbacks of the proposed approach. Previous works showed that at higher magnetic fields (>0.25 T) T2* MRR can alleviate this aspect and reliably identify brain hemorrhages, see for example [34].

The adoption of hierarchical clustering in this study, is supported by the fact that this approach has been previously applied in the histopathological field to distinguish different tissue features, see for example [36–38] and lesions characterized by different patterns [39–41]. In the case of subtle lesions, such identification with standard MRI techniques can be very and any advanced analysis tool must be validated. To this aim, here, the classification performance was directly addressed through the comparison with histopathological analyses.

As far as it strictly concerns the histopathology findings, in acute human spontaneous intracerebral hemorrhages, as well as in those induced by injecting blood in different animal models (42), primary tissue damage is followed by secondary pathological lesions, in which cellular and neuroinflammatory changes are poorly defined. Most of the studies published on this topic refer to 24/48 hours postmortem histological changes [43,44], especially focusing on the time and mass effect of the lesion, cerebral edema, and ischemic cell changes affecting the adjacent neural tissue. Consistently, in all our cases the histopathological examination on fixed tissue, one hour after suppression, showed initial alterations such as peri-hemorrhage edema as well as mild vascular and neuronal degeneration. The findings were crucial to confirm our MR tissue classification. In fact, the lesion contours extracted using the gross anatomy and histopathological images showed a good performance of the MRR classification.

However, in some cases, the adopted approach partially failed to identify the lesion. This could be to a small amount of blood that, because of heparinization, slightly infiltrated the neural tissues around the line of needle insertion during the injection. Such a small blood infiltration in the surrounding healthy parenchyma could make the MRR clustering unable to distinguish the boundary real hemorrhage-healthy tissues that instead appears evident from the histopathology after the brain fixation. This hypothesis related to heparinization is supported by the work of Dai et al. (2018), where authors, contrary to what has been herein described, used autologous nonheparinized blood into cerebral parenchyma for their experiment and observed in their results a clear distinction between hemorrhagic tissue and parenchyma [42].

In general, caution must be taken in comparing the histopathological and MRI analyses, for several reasons, e.g different slice thickness on MRI images and gross anatomy, potential misalignment, the effect of formalin fixation. Specifically, it has been reported that there is a reduction in tissue volume of approximately 33% after formalin fixation and paraffin embedding [45]. This may have influenced the data comparison, since after tissue fixation the size of the original lesion investigated with relaxometry may have changed, thus relying on a possible bias during results data co-registration and agreement quantification. Nevertheless, the obtained sensitivity and specificity place the classifier at the edge between moderate and highly accurate.

The current results pave the way for future Artificial Intelligence (AI) based techniques which are being developed to assist physicians in improving their diagnosis [46]. In this scenario, after being trained, a classification algorithm automatically maps the observations to a set of classes. Our work can be considered seminal for future AI applications validating a combined T1/T2 MRR hierarchical clustering for the classification step [47,48]. At the current stage, in our approach, the training is missing, and the proposed procedure is semi-automatic relying on a manual selection of candidate ROIs. However, the algorithm training requires datasets whose size is larger than our current sample. Therefore, it represents a future development of this technique where part of the data will train the classifier, and the remaining will be automatically classified [49,50].

To conclude, to the best of our knowledge, this is the first report on an advanced MRR classification tested on a routinely low field scanner showing that MRI diagnostic potential could be improved especially in animal brain disorders causing subtle lesions (such as e.g distemper encephalitis or hepatic encephalopathy).

## Acknowledgments

The authors would like to acknowledge the contributions of Giovanni Angelozzi (for having built the guillotine), and Marina Baffoni (for technical assistance).

## Supporting Information

**S1 Fig Analysis pipeline**

A) The voxels manually selected on the suspected lesion (black dots) are automatically extracted on the contralateral hemisphere (red dots) based on the brain midline. This step is based on the T2 map.

B) T1 based signals are extracted from the selected set of voxels. These patterns will be input to the hierarchical classification.

C) The final dendrogram for the data is pointed, and it can be seen that the cluster silhouette suggested that optimal separation corresponded to two clusters respectively healthy and hemorrhagic tissue.

D) The classifier is validated through the histopathological results. The validation is based on the true lesion contour and quantified employing ROC parameters such as TPR, FPR, and accuracy.

